# DNA damage alters nuclear mechanics through chromatin reorganisation

**DOI:** 10.1101/2020.07.10.197517

**Authors:** Ália dos Santos, Alexander W. Cook, Rosemarie E Gough, Martin Schilling, Nora Aleida Olszok, Ian Brown, Lin Wang, Jesse Aaron, Marisa L. Martin-Fernandez, Florian Rehfeldt, Christopher P. Toseland

## Abstract

DNA double-strand breaks (DSBs) drive genomic instability. For efficient and accurate repair of these DNA lesions, the cell activates DNA damage repair pathways. However, it remains unknown how these processes may affect the biomechanical properties of the nucleus and what role nuclear mechanics play in DNA damage and repair efficiency.

Here, we used Atomic Force Microscopy (AFM) to investigate nuclear mechanical changes, arising from externally induced DNA damage. We found that nuclear stiffness is significantly reduced after cisplatin treatment, as a consequence of DNA damage signalling. This softening was linked to global chromatin decondensation, which improves molecular diffusion within the organelle. We propose that this can increase recruitment for repair factors. Interestingly, we also found that reduction of nuclear tension, through cytoskeletal relaxation, has a protective role to the cell and reduces accumulation of DNA damage. Overall, these changes protect against further genomic instability and promote DNA repair. We propose that these processes may underpin the development of drug resistance.

## INTRODUCTION

Cells are known to respond to external stimuli through changes to their biomechanical properties, such as cellular stiffness. Some studies have previously related chemotherapy treatments to changes in the stiffness of cells and tissues^1,2^. Chemotherapy agents such as cisplatin induce DNA damage, and therefore their main mechanism of action occurs within a nuclear context. However, little is known about the biomechanical changes that might occur in the organelle following DNA damage.

Cisplatin, specifically, creates adducts within the double helix, which then lead to double-strand breaks (DSBs) in the DNA during replication, through replication-fork collapse^3^.

DSBs can result in large genomic aberrations and are, therefore, the most deleterious to the cell. In the event of a DNA break, the cell activates DNA Damage Response (DDR) pathways that allow detection and repair of this lesion. Failure to repair damage leads to cell death through apoptosis, or to the propagation of mutations that drive genomic instability and cancer development 4-6

It is well known that the ataxia-telangiectasia mutated (ATM) kinase localises to regions of damage, where it phosphorylates histone H2AX, producing γH2AX. This, in turn, promotes the recruitment of multiple repair factors to foci of damage^7,8^. However, it is not yet known if DNA damage, or DDR itself, could lead to alterations in nuclear mechanics.

It is established that chromatin and the lamina are major determinants of nuclear mechanics^9^, and we have previously shown that a thicker nuclear lamina correlate with higher nuclear stiffness^10^. In the context of DDR, local changes to the condensation state of the chromatin are associated to efficiency of DNA repair, and chromatin remodelling factors are known to be recruited to foci of damage^11,12^. As with DNA damage, it is still unknown how these alterations impact the physical properties of the nucleus. Here, we investigated the relationship between DNA damage and nuclear mechanics. We used Atomic Force Microscopy (AFM) to probe mechanics of the nucleus of mammalian cells and monitor changes that occur after treatment with cisplatin. We found that, following DNA damage, large-scale mechanical alterations to the nucleus arise, caused by global decondensation of chromatin. This is dependent on DDR activation and increases molecular diffusion in the nucleus, potentially resulting in higher accessibility for repair factors. Surprisingly, we also found that mechanical relaxation of the nucleus, independently of chromatin, protects against genomic instability. Collectively, our data reveal how changes to chromatin architecture following DNA damage both suppress further genomic instability and provide an environment to promote repair. These findings may be harnessed to promote therapeutic approaches *(e.g.* resistance within a tumour environment).

## RESULTS

Whilst the biochemical responses to DNA damage are well investigated^4,13,14^, their effects on the biophysical properties of the nucleus are still poorly understood. To investigate this, we used cisplatin, a chemotherapy drug that crosslinks DNA, thereby inducing DSBs within DNA, and AFM to measure mechanical changes in nucleus.

### Cisplatin treatment reduces nuclear stiffness

Before AFM measurements, fully-adhered cells on glass were treated with 25 μM cisplatin for 4 hours, or 24 hours, which induced DNA damage, recorded as fluorescent foci of γH2AX (Fig. 1A). AFM measurements were performed at a central point above the nucleus of a selected cell, up to a maximal force of 10 nN (Fig. 1B and C). Force-distance curves were then fitted using the Hertz model, to determine the effective Young’s elastic moduli *E.* Figure 1D shows representative experimental curves of deflection versus probe-sample separation and, in the insets, the fitted curves.

**Figure 1.**
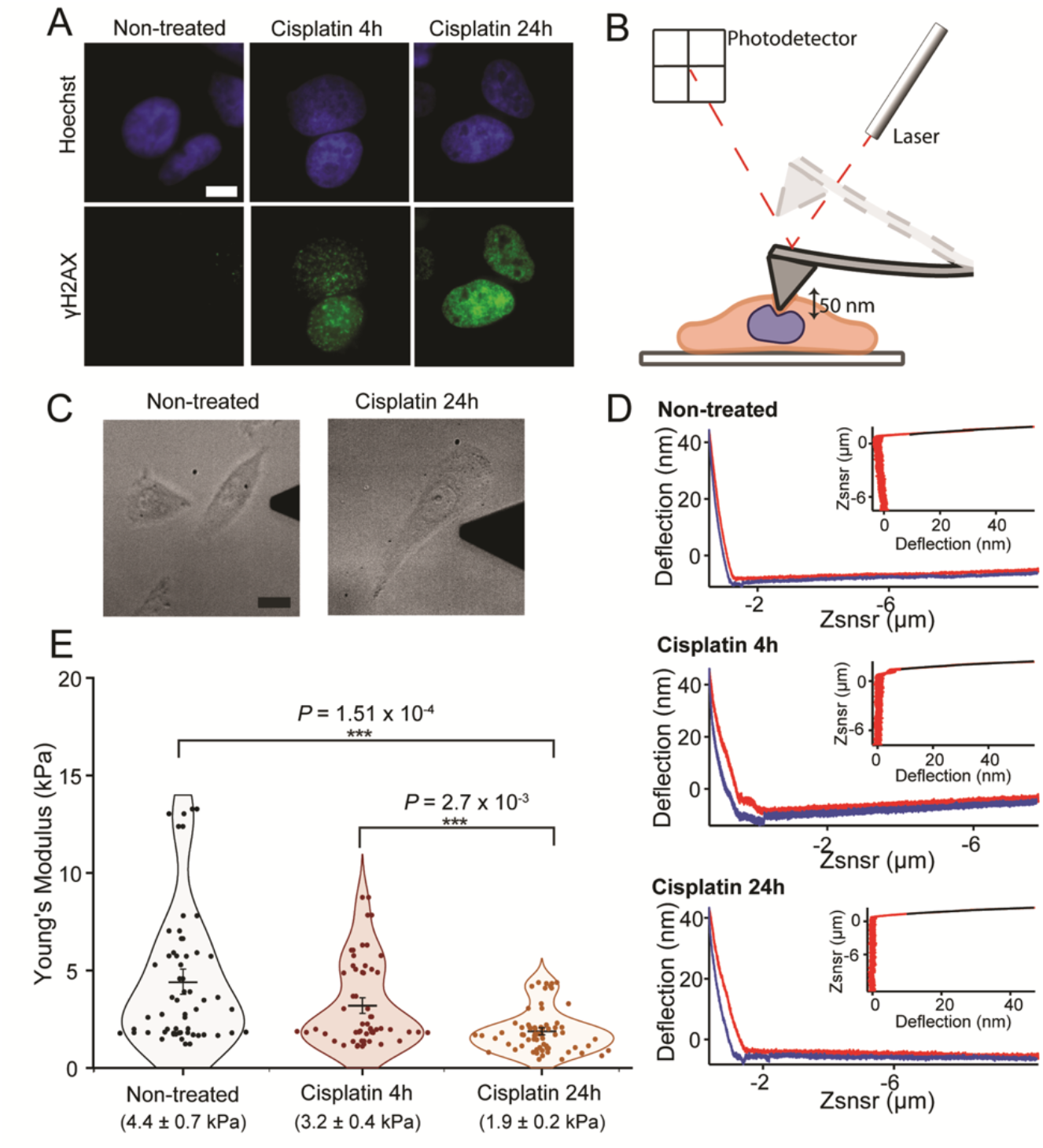
AFM measurement of cisplatin treated cells. **(A)** Wide-Field immunofluorescence imaging of γH2AX (green) in HeLa cells (scale bar = 10μm). **(B)** Cartoon depicting AFM measurement. Cells are attached on glass, and a cantilever with a pyramidal tip is used to probe cellular mechanics at a central point above the nucleus. **(C)** Transmitted light sample images from wide-field microscope coupled to AFM, showing non-treated cells and cells after long cisplatin treatment (24 hours). Scale bar 20 μm. **(D)** Representative distance (Zsnsr) versus deflection AFM curves for all three conditions tested are shown, with approach and retraction curves in red and blue, respectively. Curve fitting, shown above curves, was performed using the Hertz model, as described in methods **(E)** Young’s moduli values for non-treated *(n= 29),* cisplatin 4h (*n=29*) and cisplatin 24h *(n=34)* treatments. Each point corresponds to the average value for a cell, calculated from ten measurements. Mean ± SE are represented in the plot and values are shown below each condition. *p-values* were calculated by a two-tailed t-test, assuming equal variance; *\*\*\*p<0.001.*

We found that long treatment with cisplatin (24 hours), but not short treatment (4 hours), significantly reduced cellular stiffness (Fig. 1E). Young’s modulus values changed from 4.4 ± 0.6 kPa (mean ± SEM) in non-treated cells to 3.2 ± 0.4 kPa after cisplatin 4-hours and 1.9 ± 0.2 kPa in cells, following 24-hour cisplatin treatment. These results suggest that the observed mechanical change is temporally separated from the DNA damage itself. Therefore, breaks in DNA do not directly change the mechanical properties of the nucleus, but rather trigger events that further downstream affect the nuclear stability. Although these measurements were taken at a central point above the nucleus, the complex actin cytoskeleton extends throughout the whole cell and exerts large amounts of force on the nucleus through compression, thus altering its physical properties and potentially masking mechanical changes. As a result, it is difficult to understand what the individual contributions of the nucleus and the cytoskeleton are to wholecell mechanics.

To understand if the mechanical changes observed were a result of alterations to the cytoskeleton, we labelled cells with phalloidin after cisplatin treatment (Supp Fig.1). Our data show that there is no actin cytoskeleton impairment, indicating that the observed mechanical effect is not a result of the dissociation of actin filaments in the cell. We therefore postulated that the change in stiffness could be the result of altered biophysical properties in the nucleus.

To understand the individual contribution of the nucleus to global mechanics, we decided to perform measurements on cells at initial adhesion stages, isolated nuclei (Fig. 2A, Supp Fig 2) and cells treated with two different cytoskeletal destabilizing drugs – Blebbistatin and Latrunculin B (Supp. Fig. 1).

**Figure 2.**
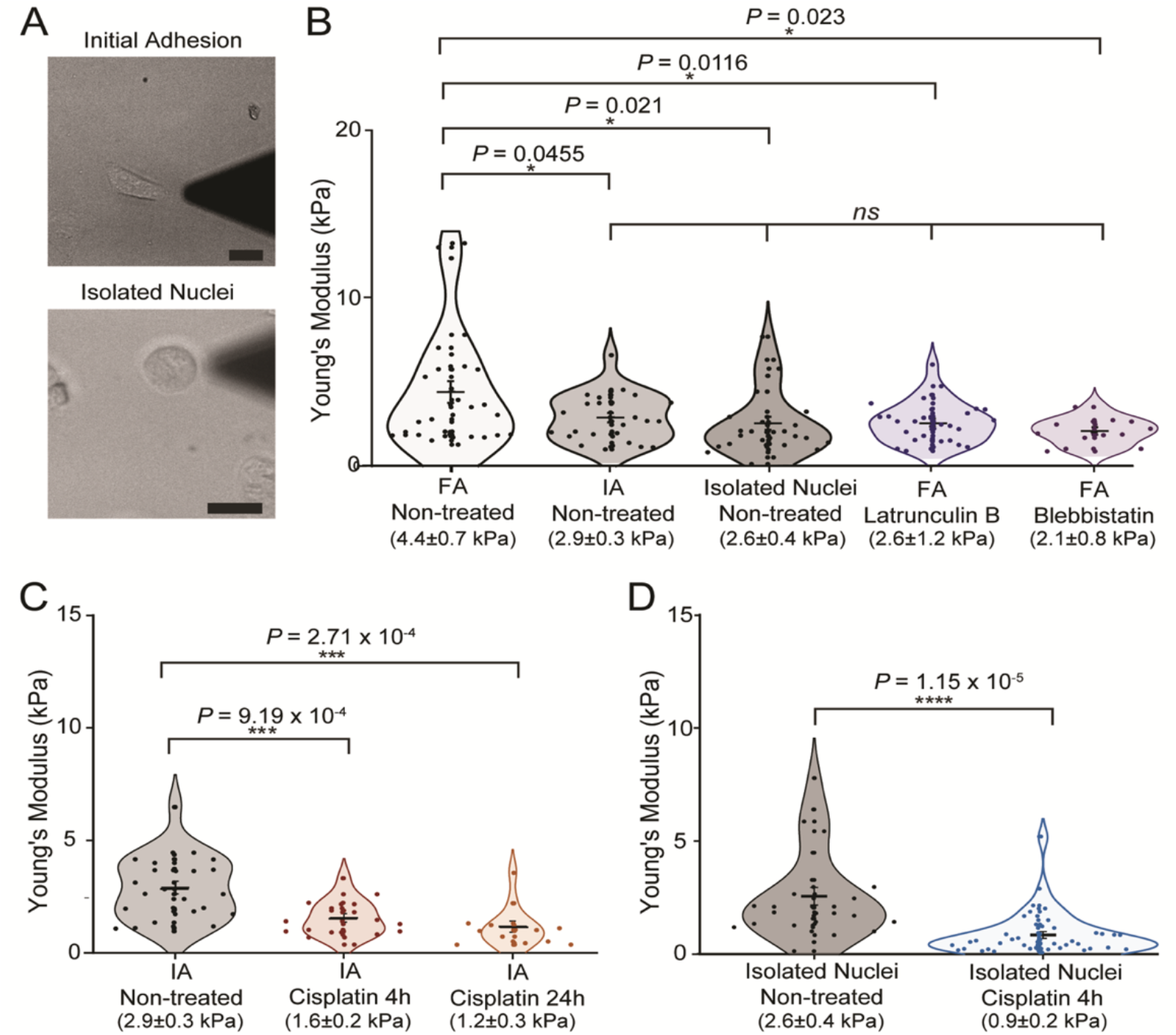
Initially adhered and isolated nuclei measurements in HeLa cells after DNA damage. **(A)** Representative transmitted light images from wide-field microscope coupled to AFM, showing initially adhered cells and isolated nuclei. (Scale bar 20 μm) **(B)** Young’s moduli values for comparison between fully adhered cells (*n=29*), initially-adhered cells (*n= 26*), isolated nuclei (*n=* 32) and after cytoskeleton disruption with drugs LatB (*n=* 28) and Blebbistatin *(n= 13).* **(C)** Young’s moduli of initially adhered cells after 4 (*n = 17)* and 24-hour (*n = 13*) cisplatin treatments. **(D)** Values for AFM measurements of non-treated isolated nuclei and nuclei isolated after 4hour cisplatin treatment (*n= 40).* Mean values ± SE are represented in the plot. *p-values* were calculated by twotailed t-test, assuming equal variance; *ns>0.05 *p<0.05; **p<0.01; ***p<0.001; ****p<0.0001.*

At initial adhesion stages, the nucleus is the largest contributor to whole-cell mechanics. In this case, the cells are given enough time to adhere to the surface (2-3 hours for HeLa cells) but the cytoskeleton and in particular stress fibres, are not fully established (Supp Fig.1). As a result, AFM measurements taken above the nucleus only reflect the mechanical properties of the organelle. Alternatively, Blebbistatin, a Myosin II inhibitor, and Latrunculin B, an actin depolymerising drug, relax the cytoskeleton^15–17^, and therefore measurements in these cells are mainly directed at the nucleus, without a contribution from the cytoskeleton.

Our AFM data show that, as expected, fully-adhered cells (4.4±0.7 kPa) are stiffer than Latrunculin B and Blebbistatin treated cells (2.6±1.2 kPa and 2.1±0.8 kPa, respectively), initially-adhered cells (2.9±0.3 kPa) and isolated nuclei (2.6±0.4 kPa), whilst there is no significant difference between these four latter conditions (Fig. 2B).

From these data we confirmed that nuclear measurements performed in initially-adhered cells are relatively free of cytoskeletal contributions and that the nucleus remains as the only large structural variable. Therefore, we decided to use this approach to investigate changes to nuclear mechanics, as this allows for the physical properties of nucleus to be probed whilst maintaining the organelle in its physiological environment and without the use of drugs that may have unknown effects in our study.

To investigate effects of DNA damage in nuclear mechanics, we used 25 μM cisplatin treatments for 4 and 24 hours on initially-adhered HeLa cells, which were seeded on glass slides 2 hours before AFM measurements. This revealed that the Young’s moduli of nuclei were significantly reduced after both 4 hours (1. 6±0.2 kPa, P<0.001) and 24 hours (1.2±0.3 kPa, P<0.001) of cisplatin treatment (Fig. 2C). Surprisingly, we could observe mechanical changes after short cisplatin treatment in initially-adhered cells (Fig. 2C), whilst this was not possible in fully-adhered (Fig. 1C). This suggests that cytoskeletal contributions, present in fully adhered cells may be masking smaller nuclear mechanical changes occurring in shorter treatments.

To confirm that the observed effect is intrinsic to the nucleus, we also performed these measurements on isolated nuclei (Fig. 2A), following 4-hour cisplatin treatment. Immunofluorescence images of isolated nuclei with membrane dye DiD show that the nuclear membrane in isolated nuclei is intact (Supp Fig. 2). As expected, there is a large decrease in Young’s moduli values from non-treated nuclei (2. 6±0.4 kPa) to cisplatin-treated (0.9±0.2 kPa, P<0.0001), confirming these changes occur are intrinsic to the nucleus (Fig. 2D).

### DNA damage signalling is required for mechanical changes to the nucleus

As cisplatin treatment causes severe DNA damage, we wondered if these mechanical changes were dependent on DDR signalling. Following DSBs, ATM kinase is recruited to sites of damage, where it phosphorylates histone H2AX^18^. This results in the recruitment of repair factors to DSBs and biochemical changes that determine the cell’s fate^7,19–21^.

To investigate this, we inhibited the ATM kinase, using inhibitor KU55933 (iATM) by pre-treating cells with iATM for 30 minutes before 4-hour treatment with 25μM cisplatin (also in the presence of iATM). Immunofluorescence shows that treatment with iATM impairs the formation of γH2AX foci (Fig. 3A). AFM measurements on initially adhered cells (Fig. 3B) show no difference between nuclei of non-treated cells (2.9±0.3 kPa) and nuclei of cells treated with cisplatin after preincubation with iATM (2.7±0.3 kPa). Surprisingly, these data suggest that mechanical changes to the nucleus occur after DNA damage response signalling is activated and appears to be dependent on ATM kinase. This confirms that mechanical changes do not arise directly from the induction of the DSBs.

**Figure 3.**
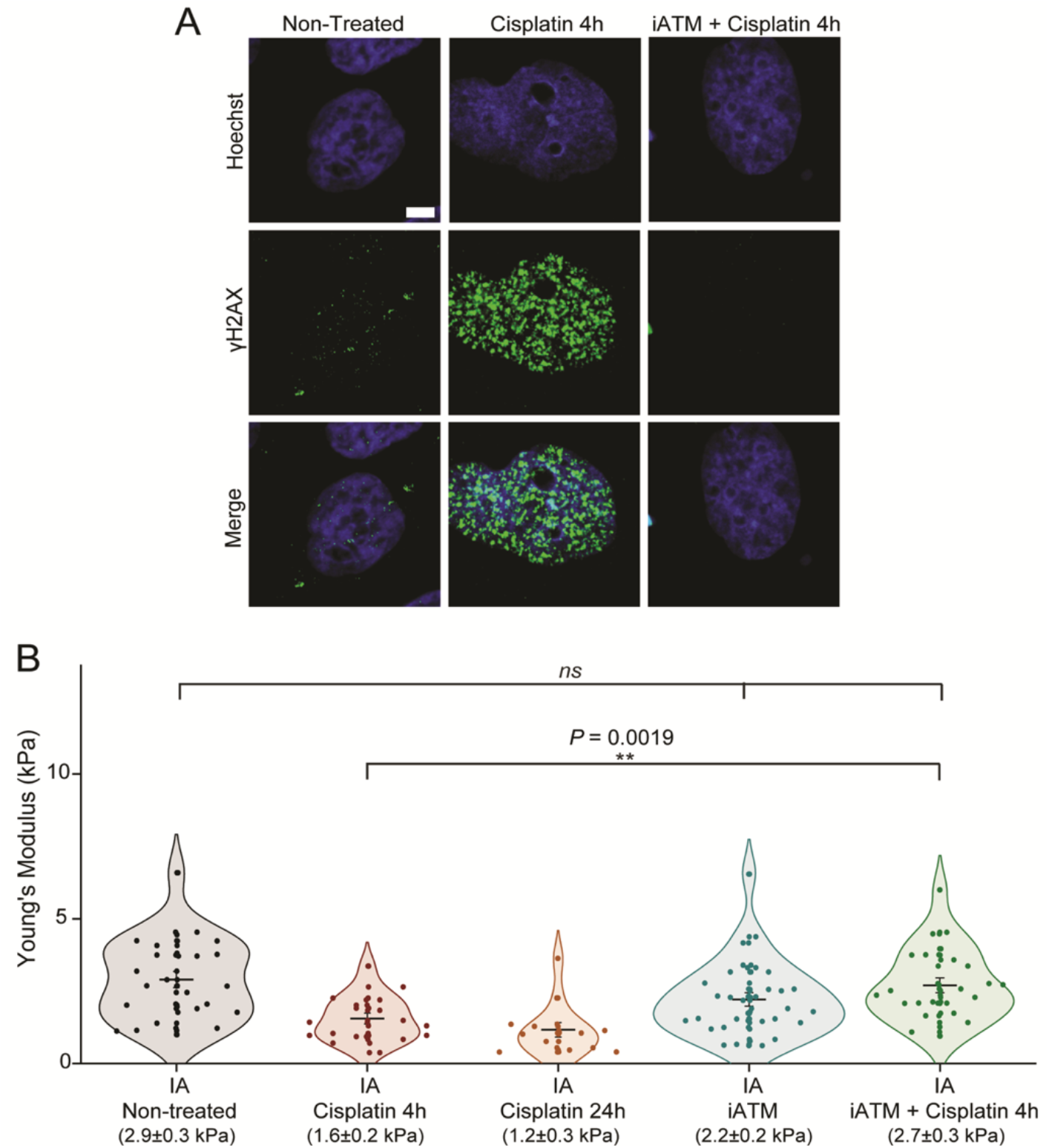
ATM inhibition impairs nuclear mechanical response to DNA damage. **(A)** Confocal immunofluorescence imaging of γH2AX (green), with nuclear stain Hoechst, after cisplatin treatment in the presence or absence of ATM inhibitor (Scale bar = 10μm). **(B)** Young’s moduli values after short and long cisplatin treatments, and in cells treated with both ATM inhibitor and cisplatin (*n = 24*). Cells treated with only iATM are also shown (*n = 31*). Plot shows mean values ± SE. *p-*values from two-tailed t-test, assuming equal variance between conditions are also shown *(ns>0.05; *p<0.05; **p<0.01).*

### Mechanical changes of the nucleus after DNA damage are caused by chromatin decondensation

The nuclear lamina is a major structural component of the nucleus. We therefore tested if cisplatin treatment changed lamina integrity, thus changing nuclear mechanics. Our results show that for non-treated and cisplatin-treated cells there are no changes in thickness of the nuclear lamina (Supp. Fig.3A and B). Negative-stain electron microscopy (EM) data support this observation and show that nuclear membrane integrity is not compromised following cisplatin treatments (Supp. Fig. 3C). This suggests that the reduction of the Young’s modulus of the nucleus is not a result of a structural compromise to the nuclear lamina.

Together with the lamina, chromatin compaction is one of the other major contributors for nuclear mechanics, and previous studies have suggested that the state of chromatin condensation can change following DSBs^12,22–24^. If this is the case, global changes to chromatin compaction could result in significant alterations to nuclear mechanics.

To investigate this hypothesis, we used EM to image both non-treated and cisplatin-treated cells and compared it to Trichostatin A (TSA), a deacetylase inhibitor that leads to decondensation of chromatin. Dense regions of staining corresponding to condensed chromatin are clearly visible in the non-treated cells. This contrasts with both cisplatin and TSA treated cells (Fig. 4A). Quantification of chromatin compaction from EM images, based on intensity of staining, shows that cells treated with cisplatin have higher levels of decondensed chromatin, similarly to TSA-treated cells, whilst non-treated cells have higher amounts of condensed chromatin (Fig. 4B).

**Figure 4.**
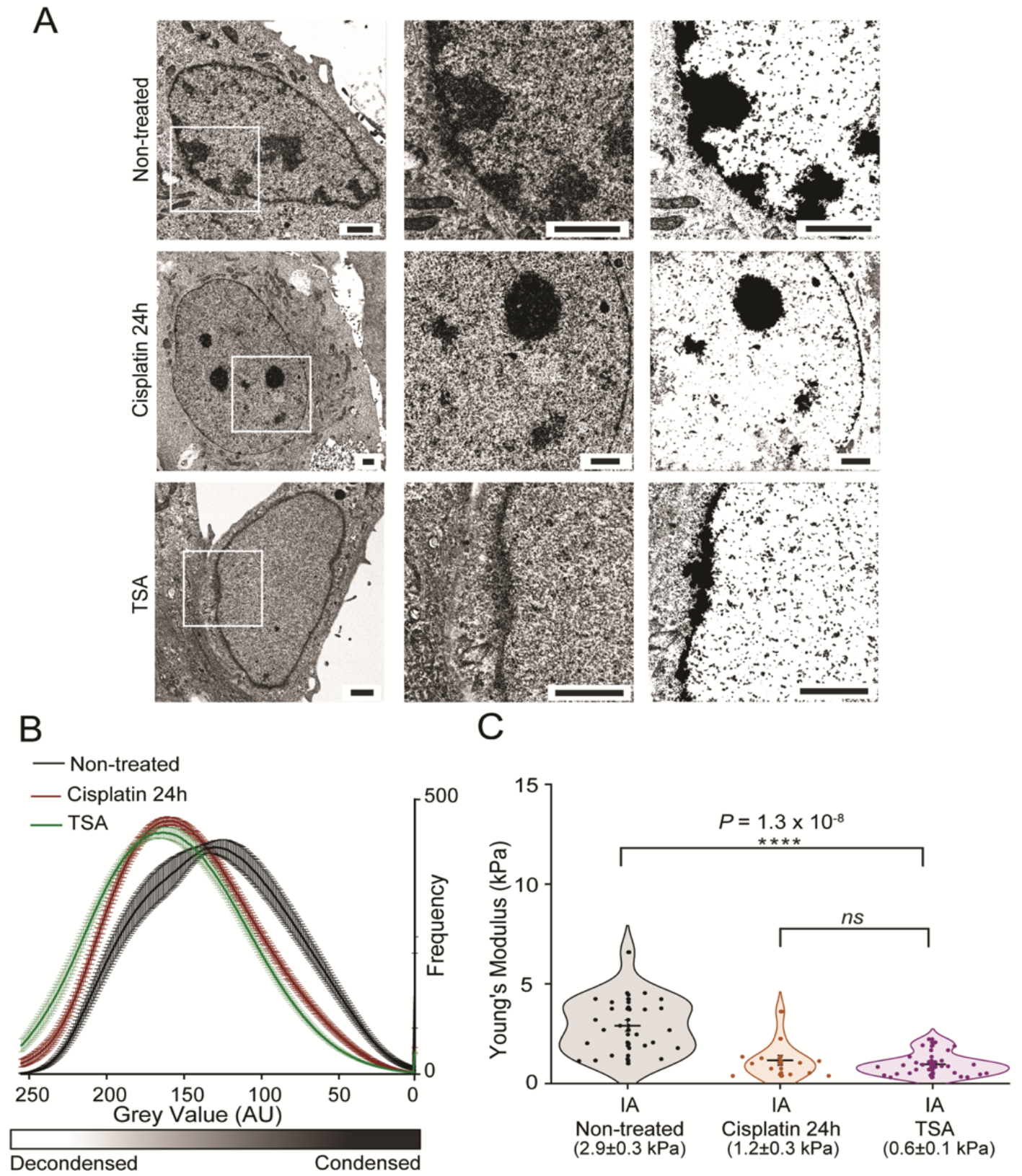
Electron Microscopy for quantification of chromatin condensation levels. **(A)** Electron microscopy of HeLa cells after long cisplatin treatment and treatment with deacetylase inhibitor TSA. White squares on left panel represent area selected for detail on middle panel. Threshold images are shown on the right panel (Scale bars = 2μm). **(B)** Quantification of dark and light pixels from electron microscopy images inside the nucleus, representing condensed and decondensed chromatin. Values for non-treated cells are in black (*n = 21*), long cisplatin treatment in red (*n = 26*) and TSA (*n = 26*). **(C)** Young’s moduli values of initially adhered cells, comparing non-treated and 24-hour cisplatin treatment with TSA (*n = 28*). For all experiments, the mean values ± SE are plotted. Statistical differences were calculated using two-tailed t-test, assuming equal variance between conditions and *p-*values are shown *(ns>0.05; *p<0.05; **p<0.01;* \*\*\**p<0.001; ****p<0.0001).*

To test if chromatin decondensation, following TSA treatment, had a similar effect to cisplatin on nuclear stiffness, we measured the Young’s modulus in nuclei of initially adhered cells. Our AFM data show (Fig. 4C) that nuclei in TSA-treated cells (1.0±0.1 kPa) are significantly softer than in non-treated cells (2.9±0.3) and display similar mechanics to long-term cisplatin-treated cells (1.2±0.3).

To complement EM data and confirm the observed effects of cisplatin on chromatin, we used super-resolution STORM imaging. Standard antibody staining was used to image γH2AX, whilst we took advantage of the photophysical properties of the Hoechst DNA dye to visualise chromatin (Fig. 5A and B). Qualitatively, our results confirm that chromatin decondensation occurs after cisplatin treatment (Fig. 5A) compared to the non-treated sample, and similarly to TSA-treated cells (Fig. 5B). In the non-treated sample, clearly defined chromatin bundles are present within the nuclear periphery and interior. However, upon cisplatin treatment there are a few islands of condensed DNA but there is loss of the extensive network. The extent of chromatin relaxation in this case is comparative to TSA treatment. Taken together, these results strongly suggest that chromatin decondensation is a determining factor for changes in nuclear mechanics after DNA damage.

**Figure 5.**
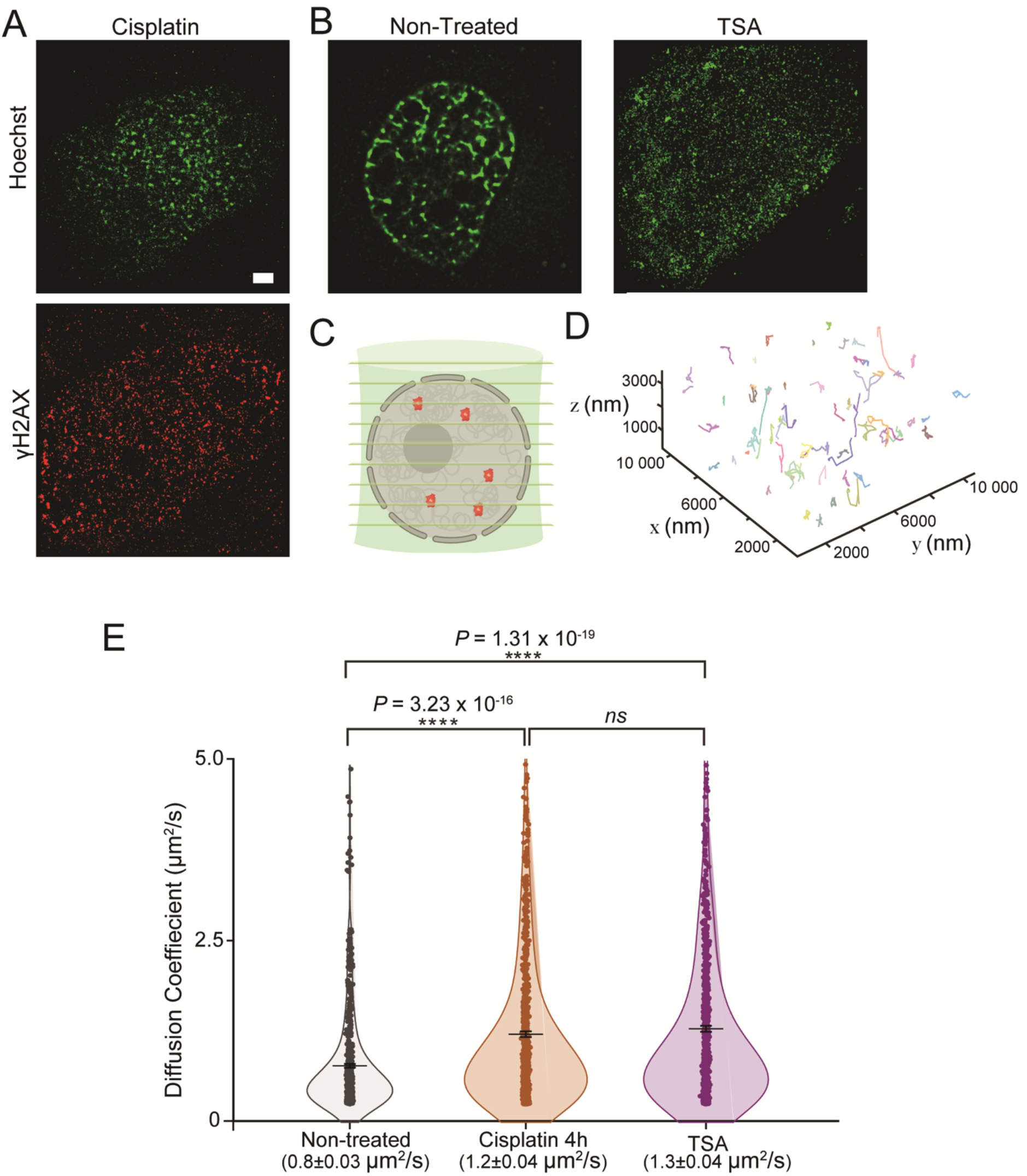
Impact of DNA damage on chromatin organisation and molecular diffusion in the nucleus. **(A)** Representative STORM images of chromatin labelled with Hoechst of HeLa cells after cisplatin treatment, with immunolabelling of γH2AX (red). Scale bar = 2 μm. **(B)** STORM images of chromatin in non-treated and TSA treated HeLa cells. **(C)** Cartoon depicting simultaneous acquisition of nine focal planes using multifocal microscopy for 3D single-molecule tracking of a reporter SNAP-tag. **(D)** Example of 3D molecule trajectories under normal conditions. **(E)** Diffusion Coefficients calculated from 3D single-molecule tracking after fitting trajectories assuming an anomalous diffusion model. Plot shows mean values ± SE. *p-*values were calculated with a two-tailed t-test, assuming equal variance *(ns>0.05; *p<0.05; **p<0.01; ***p<0.001; ****p<0.0001).*

A possible benefit that arises from chromatin decondensation is that it may allow for higher protein diffusion in the nucleus, thus enabling repair factors to reach areas of damage more easily. To test this, we used Multifocal Microscopy (MFM)^25^ and 3D single-particle tracking to measure the diffusion constants of a fluorescently-labelled SNAP tag^26,27^, expressed in HeLa cells, in treated and non-treated conditions. MFM allows us to do live-cell, singlemolecule imaging across nine simultaneous z planes (Fig. 5C)^25^, hence creating a comprehensive 3D model of molecular diffusion in nucleus. The SNAP tag was used as a reporter of free diffusion within the nucleus. Figure 5D shows a representative 3D map for single-particle tracking using MFM.

Our data show that the diffusion constant of the SNAP-tag in the nucleus of cells treated with cisplatin (1.2±0.04 μm^2^/s) is significantly higher than in non-treated cells (0.8±0.03 μm^2^/s), but similar to TSA-treated (1.3±0.04 μm^2^/s) (Fig. 5E). This supports the idea that chromatin decondensation following damage allows proteins to diffuse more rapidly within the nucleus.

### Mechanical relaxation of the nucleus protects cells from damage

Recently, studies have linked the occurrence of DNA damage to external forces exerted on the nucleus^28,29^. Similarly, there is some evidence that increased forces on the nucleus exacerbate the extent of DNA damage^30,31^. It is possible that, after DNA damage, a softer nucleus, also leads to a loss of tension on the nucleus which prevents further genomic instability.

To investigate how forces exerted on the nucleus influence DNA damage, we decided to pre-treat cells with Blebbistatin for 30 minutes before 4-hour treatment with cisplatin (also in the presence of Blebbistatin). This allowed us to minimise cytoskeletal forces on the nucleus prior to the occurrence of damage. We used γH2AX as a marker for damage. High-content screening revealed a large decrease in the percentage of damaged cells when they pre-incubated with Blebbistatin. In non-treated cells, only 5.1±0.2% of the population showed γH2AX signalling, as expected. 37.4±0.9% of cisplatin-treated cells (short treatment) displayed signs of damage, whilst this value dropped to 21.2±0.6% for cells that were pre-incubated with Blebbistatin (Fig. 6A and B). This suggests that a decrease in forces acting on the nucleus has a protective effect towards DNA damage.

**Figure 6.**
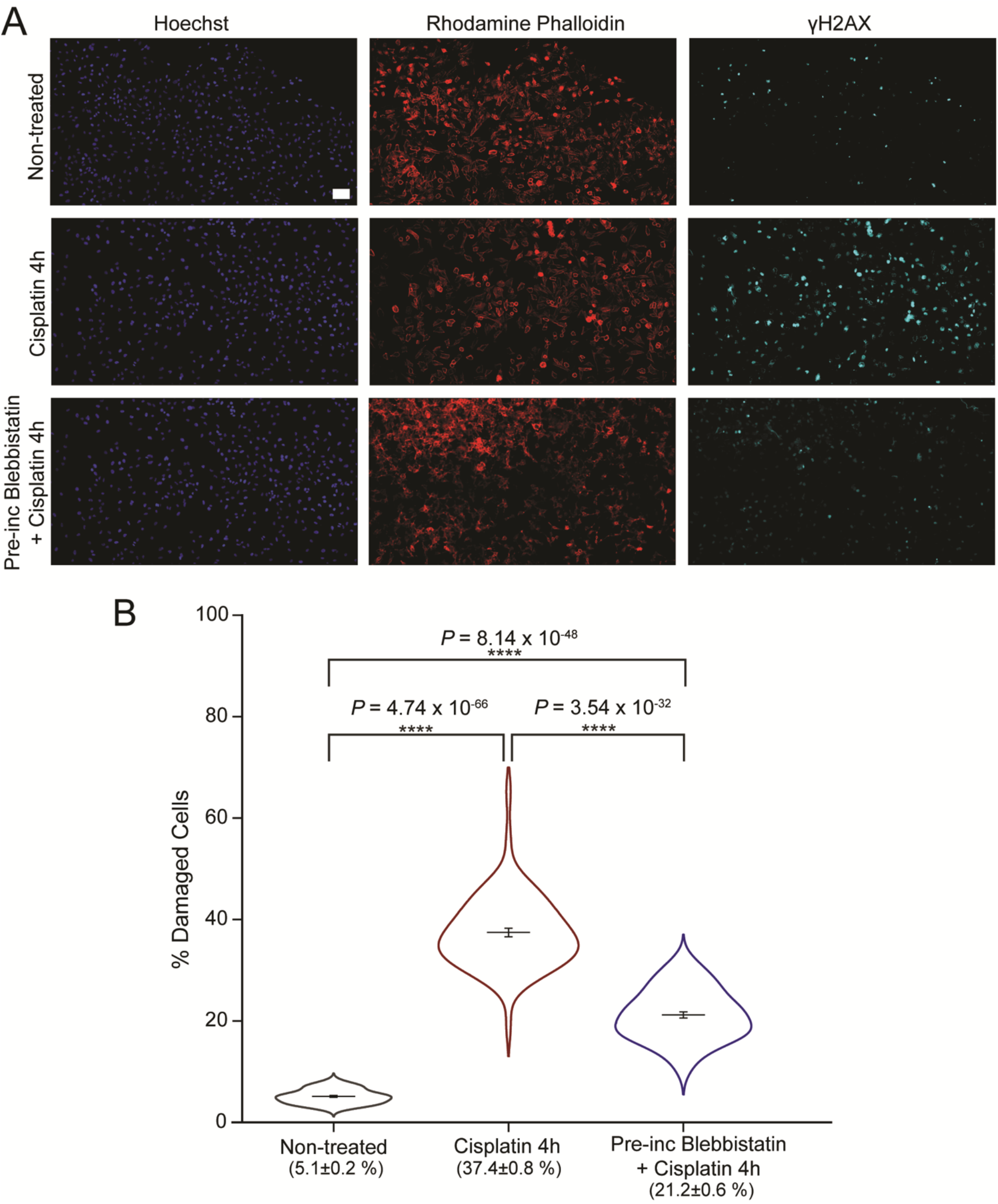
Impact of Blebbistatin on γH2AX signalling in HeLa cells. **(A)** Representative images from high-content screening of HeLa cells treated with cisplatin in the presence or absence of Blebbistatin. Nuclear staining with Hoechst and Actin labelling with Rhodamine phalloidin are shown in addition to immunofluorescent labelling of γH2AX. Scale bar = 100μm. **(B)** Levels of damage in HeLa cells from high-content screening, calculated as percentage of nuclei displaying γH2AX signalling. *P* values are shown between non-treated cells (*n = 58*, representing 51 233 cells), 4-hour cisplatin treatment (*n =70*, representing 63 422 cells) and combined Blebbistatin and cisplatin treatment (*n = 68*, representing 57 274 cells). For all conditions, mean values ± SE are plotted. Two-tailed t-test, assuming equal variance, was used for *p-value* calculation *(ns>0.05; *p<0.05; **p<0.01; ***p<0.001; ****p<0.0001).*

Matrix stiffness is tightly related to cell spread, adhesion formation and cytoskeletal force^32–34^. As a result, a stiffer surface will create higher forces in the cytoskeleton, which would then translate into higher mechanical constraints to the nucleus^35–37^. To further explore this relationship between stiffness and the extent of DNA damage, we used polyacrylamide gels of different stiffness (2, 11 and 30 kPa) as surfaces for cells (Fig. 7A). The ability of the cells to respond to the gel stiffness is confirmed when we measure the cross-sectional area of the nucleus. Here, the greater tension exerted by the stiffer gel matrix leads to an increase in nuclear area from 291.0± 8.5 μm^2^ to 456.0 ± 10.8 μm^2^ (Fig. 7B), in line with earlier findings^38^.

**Figure 7.**
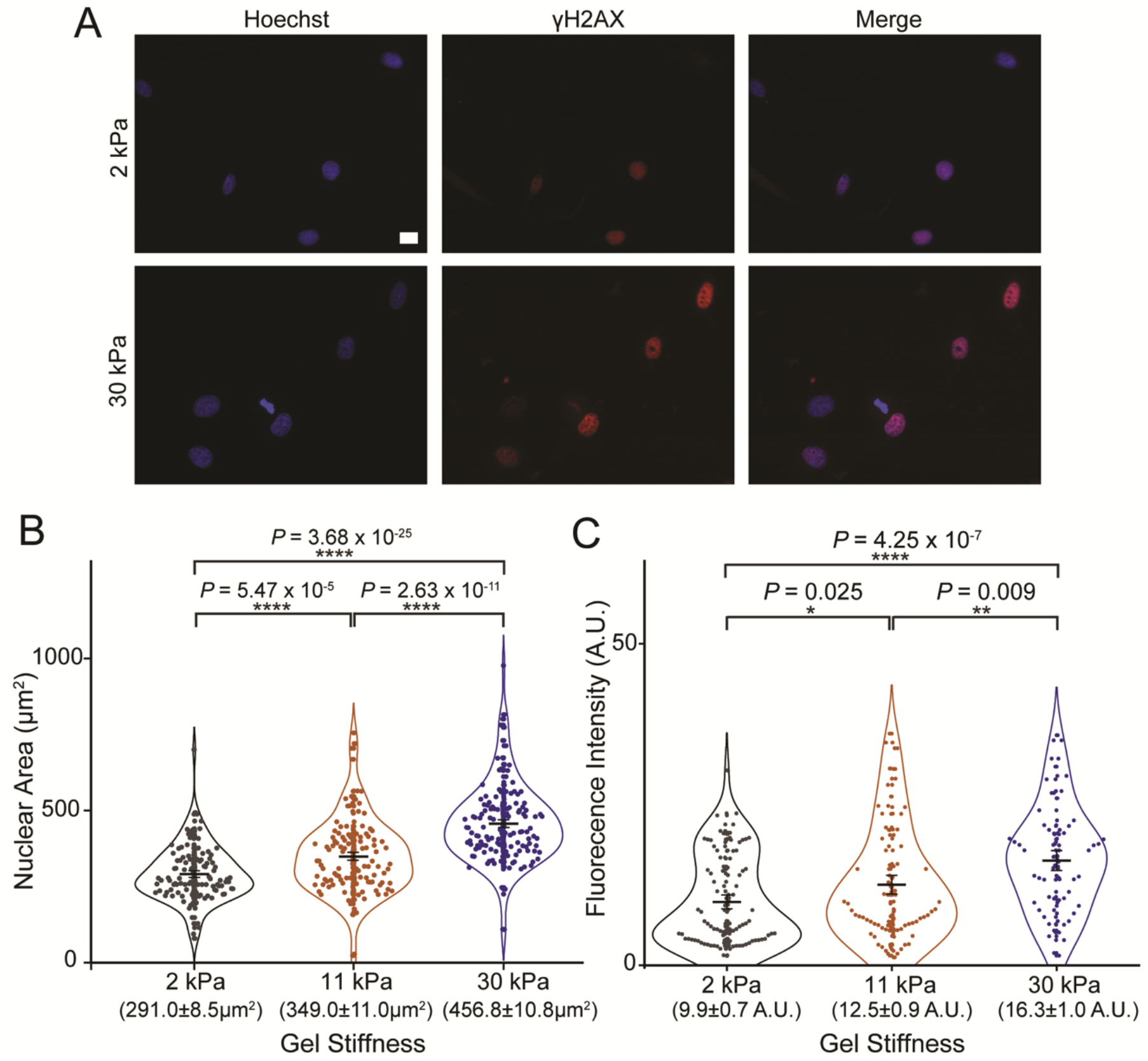
Impact of nuclear relaxation on γH2AX signalling in HeLa cells. **(A)** Representative Wide-field images of HeLa cells growing on gels of different stiffness – 2 and 30kPa – after cisplatin treatment. Immunofluorescent labelling of γH2AX is shown in red, with nuclear stain Hoechst in blue. (Scale bar = 20μm). **(B)** Nuclear area of cells on different surfaces – 2kPa *(n = 111),* 11 kPa (*n = 121)* and 30kPa (*n = 139). P* values are shown. **(C)** Fluorescence intensity of red channel (γH2AX) within the nucleus of cells grown on surfaces of different stiffness – 2kPa (*n = 90*), 11kPa (*n = 79*) and 30kPa (*n = 61*). In all cases, mean values ± SE are shown. Twotailed t-test, assuming equal variance, was used for statistical calculations and *p-*values are shown *(ns>0.05; *p<0.05; **p<0.01; ***p<0.001; ****p<0.0001).*

If we expect lower force to have a protective effect, lower surface stiffness should decrease the amount of damage in cells, in a similar way to Blebbistatin treatment. As expected, our data reveal that cells treated in stiffer gels, 30 kPa, have 65% higher levels of γH2AX levels compared to 2kPa gels and 30% higher than 11 kPa gels (Fig. 7C). Overall, less tension across the nucleus leads to less damage.

## DISCUSSION

Our work reveals a direct link between DNA damage, chromatin condensation and nuclear mechanics. It has been shown that treatment with cisplatin and other chemotherapy drugs changes whole-cell mechanics^39,40^. However, since most studies focus on cytoskeletal contributions, until now it remained unclear how these DNA damage-causing drugs alter the biomechanical properties of the nucleus. Here, by performing AFM measurements in initially adhered cells, we were able to probe the mechanical properties of the nucleus within its cellular environment. Our work shows that cisplatin treatment causes significant alterations to the state of chromatin condensation, which in turn results in a reduction of nuclear stiffness.

Mechanical changes to the nucleus do not arise spontaneously from breaks in the chromatin fibre, but are a result of downstream large-scale chromatin decondensation. The ATM kinase is an essential DSB repair factor that initiates DDR signalling, and its absence or inactivity can lead to a proportion of unrepaired DSBs in the cell^41,42^. Interestingly, the mechanical softening of the nucleus also appears to be coupled to the activity of the ATM kinase, suggesting that chromatin decondensation occurs as a result of the DDR. Several studies have reported localised chromatin decondensation around areas of damage and this is thought to lead to higher repair efficiency^22,23,43^. Our data largely agree with this observation, but we cannot exclude that there are patches of condensed DNA around the damage sites. Moreover, we show that chromatin decondensation, following DNA damage, increases molecular diffusion within the nucleus. By creating greater exposure to DNA binding sites and enabling repair factors to readily reach sites of damage, chromatin decondensation could act as a major determinant for the outcome of repair events.

Interestingly, we found that alterations to biomechanics also appear to protect the cell from genomic instability. In addition to the intrinsic mechanical properties of the nucleus, the cytoskeleton also has a large impact on overall nuclear mechanics. Relaxation of the cytoskeleton results in reduced nuclear tension and, here, we show that this has a protective effect in DNA damage. It has been shown that reduction in matrix stiffness, a major determining factor for cytoskeletal organisation, correlates with inhibition of replication^44–46^. Cisplatin-induced DSBs require active DNA replication. It is, therefore, possible that the lower levels of DSBs detected in nuclei under lower mechanical tension are a result of altered DNA replication levels. We suggest that a decrease in nuclear stiffness would also destabilise cytoskeleton-nuclear linkage which could result in a change in nuclear tension. Our findings agree with the recent findings of Nava et al, which showed that nuclear softening protects against mechanically induced DNA damage^30^. Overall, it appears that the nucleus has an innate ability to protect against different agents.

Deformations to the nuclear envelope, as well as mechanical forces acting on the organelle are important for the regulation of cell cycle progression^47^ and transcription activity^48^. Interestingly, DNA damage leads to cell cycle arrest and transcriptional repression to promote repair. Therefore, DNA damage-induced changes to the mechanosensing properties of the nucleus could contribute to the regulation of these processes. In terms of cell cancer, during migration and invasion, altered nuclear mechanics may be an important factor. For example, whilst travelling through confined spaces, cells with softer nuclei could migrate more easily and with less DNA damage, induced by rupture events resulting from nuclear compression. In this case, a softer nucleus would, once again, protect from further damage. There is growing evidence that tumorigenesis and resistance to chemotherapy agents correlate with changes in cellular and nuclear mechanics^49^. Depending on the cell types and type of treatment, drug resistance has been associated with an increase or decrease in cell stiffness^49,50^. Interestingly, there are also possible differences between *in vitro* cell lines and patient-derived primary samples, where the latter are more deformable. Our data fit with these scenarios, whereby decreased nuclear stiffness promotes repair and prevents further damage, which could subsequently drive drug resistance.

In summary, here, we describe how nuclear mechanics change due to induced DNA damage through chromatin remodelling events, as part of DDR. Furthermore, we show that nuclear envelope relaxation protects against further damage. It will be of interest, in the future, to determine the molecular mechanisms through which nuclear mechanics directly influence DNA damage levels as these pathways may directly influence therapeutic resistance.

## ACKNOWLEDGEMENTS

We thank the UKRI-MRC (MR/M020606/1), UKRI-STFC (19130001) and the Royal Society (IE170270) for funding to C.P.T. Aberration-corrected multi-focal microscopy was performed in collaboration with the Advanced Imaging Center at Janelia Research Campus, a facility jointly supported by the Howard Hughes Medical Institute and the Gordon and Betty Moore Foundation. We also thank Satya Khuon (Janelia Research Campus) for assisting with cell culture and Thomas Helleday for sharing equipment. The JF549 dye was kindly provided by Luke Lavis (Janelia Research Campus).

## AUTHOR CONTRIBUTIONS

C.P.T. and F.R. conceived the study. A.dS., F.R. and C.P.T. designed experiments. A.dS. performed AFM measurements with support from M.S. and N.A.O. A.W.C. and R.E.G. performed experiments using the acrylamide gels. I.B. prepared and collected electron microscopy data. A.dS. and C.P.T. performed single molecule imaging experiments. Imaging was supported by L.W., M.L.M-F. and J.A. L.W., J.A., A.dS. and C.P.T. contributed to single molecule data analysis. C.P.T. and F.R. supervised the study. A.dS., C.P.T. and F.R. wrote the manuscript with comments from all authors.

## MATERIALS AND METHODS

### Drug Treatments

Cisplatin [*cis*-diammineplatinum(II) dichloride] (Sigma) was resuspended in a 0.9% NaCl solution to a concentration of 3.3 mM, following manufacturer’s instructions, and used at a concentration of 25μM.Trichostatin A (Sigma) was resuspended to 6.6. mM in DMSO and used at 400nM in culturing medium, for 24 hours. ATM inhibitor, KU55933 (Sigma), was resuspended in DMSO to 12.6 mM and used at a concentration of 20μM, 30 minutes prior treatment and then for the whole duration of cisplatin treatment in culturing medium. Blebbistatin (Sigma) was resuspended to 50 mM in DMSO and used at a concentration of 50μM for 1-2 hours before AFM measurements and for 30 minutes, prior treatment and then the whole duration of cisplatin treatment for imaging. Latrunculin B (Sigma) was resuspended in DMSO to a concentration of 12.6 mM and used at 1μM for 1 hour before AFM measurements.

### Cell Culture and Transfection

HeLa cells (ECACC 93021013) were cultured at 37°C and 5% CO_2_ in MEM Alpha medium with GlutaMAX, supplemented with 10% fetal bovine serum (Gibco) and 1% penicillin/streptomycin (Gibco). HeLa cells were transfected with 0.5 μg pSNAPf-C1 plasmid (Addgene 58186) using Lipofectamine 2000 (Invitrogen) and following manufacturer’s instructions.

### Nuclear Isolation

Nuclei were prepared based on protocols in^10,51^ HeLa cells were washed with cold PBS, then washed in cold Hypotonic Buffer N – 10 mM Hepes pH 7.5, 2 mM MgCl_2_, 25 mM KCl, 1 mM PMSF, 1 mM DTT and Protease Inhibitor Cocktail (Thermo Fisher Scientific). Cells were then re-suspended in cold hypotonic buffer N and incubated for 1 hour on ice. Cells were homogenised on ice, using a glass Dounce homogeniser (Wheaton). Sucrose was added to the cell lysate to a final concentration of 220 mM and mixed well by inversion before centrifugation. The pellet, corresponding to isolated nuclei, was washed in cold Buffer N – 10 mM Hepes pH 7.5, 2 mM MgCl_2_, 25 mM KCl, 250 mM sucrose, 1 mM PMSF, 1 mM DTT Protease Inhibitor Cocktail. The nuclei pellet was resuspended in PBS and used immediately for AFM measurements.

### Immunofluorescence

HeLa cells grown on glass coverslips were incubated for 10 minutes at 37°C and 5% CO_2_ with 1μg mL^-1^ Hoechst 33342 in growth medium. Stained cells were fixed in 4% (m/V) paraformaldehyde (PFA). Residual PFA was quenched with 50 mM ammonium chloride for 15 minutes at room temperature. Cells were then permeabilised and blocked for 15 minutes with 0.1% (V/V) Triton X-100 and 2% (m/V) BSA in TBS. Antibodies were used as follows: mouse-Phospho-H2AX (Merck 05-636) at 1:500 dilution, rabbit-Lamin B1 (Abcam ab16048) at a dilution of 1:200, donkey anti-mouse Alexa Fluor 488-conjugated (Abcam Ab181289) at 1:500 and donkey anti-rabbit Alexa Fluor-488-conjugated (Abcam, Ab81346) at 1:500 dilution. For actin staining, fixed and permeabilised cells were stained prior to immunofluorescence with 165 nM Rhodamine-Phalloidin (ThermoFisher) for 20 min. Coverslips were mounted on microscope slides with 10% (m/V) Mowiol, 25% (m/V) glycerol, 0.2 M Tris-HCl, pH 8.5, supplemented with 2.5% (m/V) of DABCO (Sigma).

### Fluorescence Imaging

Cells were visualised using Wide-field microscope Olympus IXT1, or Confocal microscope LSM 880. For confocal microscopy, a Plan-Apochromat 63x 1.4 NA oil immersion lens (Carl Zeiss, 420782-9900-000) was used. Three laser lines: 405 nm, 488 nm and 561 nm, were used to excite Hoechst, Alexa 488 and Alexa 647 fluorophores, respectively. Built-in dichroic mirrors (Carl Zeiss, MBS-405, MBS-488 and MBS-561) were used to reflect the excitation laser beams onto cell samples. For fluorescence collection, the used emission spectral bands were: 410 nm-524 nm (Hoechst), 493 nm-578 nm (Alexa 488) and 564 nm-697 nm (Alexa 647). The green channel (Alexa 488) was imaged using a 1 gallium arsenide phosphide (GaAsP) detector, while the blue (Hoechst) and red (Alexa 647) channels were imaged using two multianode photomultiplier tubes (MA-PMTs). For imaging acquisition and rendering, ZEN software was used. Confocal Images were deconvolved using the Zeiss Zen2.3 Blue software, using the regularised inverse filter method.

For Wide-field microscopy, a PlanApo 100xOTIRFM-SP 1.49 NA lens mounted on a PIFOC z-axis focus drive (Physik Instrumente, Karlsruhe, Germany) was used, with an automated 300W Xenon light source (Sutter, Novato, CA) with appropriate filters (Chroma, Bellows Falls, VT). QuantEM (Photometrics) EMCCD camera, controlled by the Metamorph software (Molecular Devices) was used for image acquisition. All images were then analysed by ImageJ.

### STORM Imaging

Cells were seeded on pre-cleaned No 1.5, 25-mm round glass coverslips, placed in 6-well cell culture dishes. Glass coverslips were cleaned by incubating them for 3 hours, in etch solution, made of 5:1:1 ratio of H_2_O: H_2_O_2_ (50 wt. % in H_2_O, stabilized, Fisher Scientific): NH_4_OH (ACS reagent, 28-30% NH_3_ basis, Sigma), placed in a 70°C water bath. Cleaned coverslips were repeatedly washed in filtered water and then ethanol, dried and used for cell seeding. Transfected or non-transfected cells were incubated with 1μg mL^-1^ Hoechst 33342 in growth medium for 15 minutes at 37°C, 5%CO_2_. Following this, cells were washed with PBS and fixed in pre-warmed 4% (w/v) PFA in PBS and residual PFA was quenched for 15 min with 50 mM ammonium chloride in PBS. Immunofluorescence (IF) was performed in filtered sterilised PBS, unless when anti-phospho antibodies were used. Then, IF was performed in filtered sterilised TBS. Cells were permeabilized and simultaneously blocked for 30 min with 3% (w/v) BSA in PBS or TBS, supplemented with 0.1 % (v/v) Triton X-100.

Permeabilized cells were incubated for 1h with the primary antibody and subsequently the appropriate fluorophore-conjugated secondary antibody, at the desired dilution in 3% (w/v) BSA, 0.1% (v/v) Triton X-100 in PBS or TBS. The antibody dilutions used were the same as for the normal IF protocol (see above), except from the secondary antibodies which were used at 1:250 dilution. Following incubation with both primary and secondary antibodies, cells were washed 3 times, for 10 min per wash, with 0.2% (w/v) BSA, 0.05% (v/v) Triton X-100 in PBS or TBS. Cells were further washed in PBS and fixed for a second time with pre-warmed 4% (w/v) PFA in PBS for 10 min. Cells were washed in PBS and stored at 4 °C, in the dark, in 0.02% NaN3 in PBS, before proceeding to STORM imaging.

Before imaging, coverslips were assembled into the Attofluor^®^ cell chambers (Invitrogen). Imaging was performed in freshly made STORM buffer consisting of 10 % (w/v) glucose, 10 mM NaCl, 50 mM Tris – pH 8.0, supplemented with 0.1 % (v/v) 2-mercaptoethanol and 0.1 % (v/v) pre-made GLOX solution which was stored at 4°C for up to a week (5.6 % (w/v) glucose oxidase and 3.4 mg/ml catalase in 50 mM NaCl, 10 mM Tris – pH 8.0). All chemicals were purchased from Sigma. Imaging was undertaken using the Zeiss Elyra PS.1 system. Illumination was from a HR Diode 642 nm (150 mW) and HR Diode 488 nm (100 mW) lasers where power density on the sample was 7-14 kW/cm^2^ and 7-12 kW/cm^2^, respectively Imaging was performed under highly inclined and laminated optical (HILO) illumination to reduce the background fluorescence with a 100x NA 1.46 oil immersion objective lens (Zeiss alpha Plan-Apochromat) with a BP 420-480/BP495-550/LP 650 filter. The final image was projected on an Andor iXon EMCCD camera with 25 msec exposure for 20000 frames for γH2AX and 60 msec for 60000 frames for Hoechst imaging.

Image processing was performed using the Zeiss Zen software. Where required, two channel images were aligned following a calibration using pre-mounted MultiSpec bead sample (Carl Zeiss, 2076-515). The channel alignment was then performed in the Zeiss Zen software using the Affine method to account for lateral, tilting and stretching between the channels. The calibration was performed during each day of measurements.

The images were then processed through our STORM analysis pipeline using the Zen software. Single molecule detection and localisation was performed using a 9-pixel mask with a signal to noise ratio of 6 in the “Peak finder” settings while applying the “Account for overlap” function. This function allows multi-object fitting to localise molecules within a dense environment. Molecules were then localised by fitting to a 2D Gaussian.

The render was then subjected to model-based cross-correlation drift correction. The final render was then generated at 10 nm/pixel and displayed in Gauss mode where each localisation is presented as a 2D gaussian with a standard deviation based on its precision.

### High-Content Screening

HeLa cells were seeded in clear-bottom, black-walled 96-wellplates at 10 000 cells/well in culturing medium and incubated overnight at 37°C, 5% CO_2_. For Blebbistatin pre-treated cells, 50 μM Blebbistatin was added prior to 4-hour incubation with 50 μM Blebbistatin and 25 μM cisplatin. For cisplatin-treated cells, only cisplatin was added at 25 μM for 4 hours. Cells were then incubated for 15 minutes with Hoechst dye, at 37°C, 5% CO_2_ and fixed using 4%(m/v) PFA. Cells were stained with 165 nM Rhodamine-Phalloidin (ThermoFisher) for 20 min and 1:500 dilution of mouse-Phospho-H2AX (Merck 05-636) antibody for 1 hour, following the immunofluorescence protocol. High-content imaging was undertaken using a Cell Discoverer 7 (Zeiss), using a Plan-Apochromat 20x 0.7 NA objective. Hoechst, Alexa 488 and Rhodamine fluorophores were excited using LED light at wavelengths 385, 470 and 567nm, respectively. ZEN software was used for image acquisition and images were analysed using Zeiss Zen2.3 Blue software.

### Multi-focal Imaging and Particle Tracking Analysis

Cells were transfected for 24 hours with 0.5μg of pSNAPf-C1 (Addgene 58186) construct with Lipofectamine 2000 (Invitrogen), according to manufacturer’s instructions. Following this, cells were treated with 25 μM cisplatin or 400nM TSA in growth medium for 24 hours.

Cells transiently expressing pSNAPf-C1 construct were labelled for 15 min with 10 nM SNAP-tag-JF549 ligand, in cell culture medium at 37°C, 5% CO_2_. Cells were washed for 3 times with warm cell culture medium and then incubated for further 30 min at 37°C, 5% CO_2_. Cells were then washed three times in pre-warmed FluoroBrite DMEM imaging medium (ThermoFisher Scientific), before proceeding to imaging.

Single molecule imaging was performed using an aberration-corrected multifocal microscope (acMFM), as described by Abrahamsson et al.^25^. Briefly, samples were imaged using 561nm laser excitation, with typical irradiance of 4-6 kW/cm^2^ at the back aperture of a Nikon 100x 1.4 NA objective. Images were relayed through a custom optical system appended to the detection path of a Nikon Ti microscope with focus stabilization. The acMFM detection path includes a diffractive multifocal grating in a conjugate pupil plane, a chromatic correction grating to reverse the effects of spectral dispersion, and a nine-faceted prism, followed by a final imaging lens.

The acMFM produces nine simultaneous, separated images, each representing successive focal planes in the sample, with ca. 20 μm field of view and nominal axial separation of ca. 400nm between them. The nine-image array is digitized via an electron multiplying charge coupled device (EMCCD) camera (iXon Du897, Andor) at up to 32ms temporal resolution, with typical durations of 30 seconds.

3D+t images of single molecules were reconstructed via a calibration procedure, implemented in Matlab (MathWorks), that calculates and accounts for (1) the inter-plane spacing, (2) affine transformation to correctly align each focal plane in the xy plane with respect to each other, and (3) slight variations in detection efficiency in each plane, typically less than ±5-15% from the mean.

Reconstructed data were then subject to preprocessing, including background subtraction, mild deconvolution (3-5 Richardson-Lucy iterations), and/or Gaussian de-noising prior to 3D particle tracking using the MOSAIC software suite^52^. Parameters were set where maximum particle displacement was 400 nm and a minimum of 10 frames was required. Tracks were reconstructed, and diffusion constants were extracted via MSD analysis^53^ using custom Matlab software assuming an anomalous diffusion model.

### Atomic Force microscopy

Atomic Force microscopy (AFM) measurements were performed with an MFP 3D BIO (Asylum Research, Oxford Instruments, Santa Barbara, USA) on top of an inverted fluorescence microscope (IX71, Nikon Instruments, Japan). The combined instrument is mounted on a Halcyonics vibration isolation table (Accurion GmbH, Göttingen, Germany) inside an acoustic enclosure (Asylum Research) to minimize noise. Force-indentation curves were executed in contact mode using the longer triangular cantilever of a TR 400 PB chip that has a pyramidal tip with an opening angle of 35°. The spring constant was determined with the built-in macro based on the thermal method and was in the range (28–31 pN nm^-1^) and therefore slightly stiffer than the nominal stiffness of 0.02 Nm^-1^.The nucleus was probed on several locations using the ForceMap macro with a maximum indentation force of 1.5 nN and the resulting force-indentation curves were analysed using a modified Hertz model within a self-written IGOR macro as described earlier^54,55^.

### Polyacrylamide gels

Elastic polyacrylamide (PA) gels were prepared as described earlier^34,55^. In brief, mixtures for the desired elasticities were prepared using solutions of 40% acrylamide (#161-0140, Bio-Rad, Munich, Germany) and 2% bis-acrylamide (#161-0142, Bio-Rad, Munich, Germany) in PBS that are stored at 4°C for a maximum time of 2 months. These mixtures were polymerized through addition of 1% (v/v) ammoniumpersulfate and 0.1 % (v/v) N,N,N,N-tetramethylethylenediamine onto freshly plasma-cleaned cover slips that were treated with 3-aminopropyltriethoxysilane (Sigma, Munich, Germany, A3648) and subsequently with a 0.05% glutaraldehyde solution (Sigma, Munich, Germany, G7651) for firm attachment of the gels. For 25 mm coverslips, 35 μL of the PA gel solution was dispensed and covered with a square superhydrophobic cover glass to equally distribute the solution. Gels were polymerized for 60 min using a plastic box to keep them in a saturated water atmosphere to avoid evaporation. For quality control the Young’s elastic modulus *E* was regularly measured with a bulk rheometer (MCR-501, Anton Paar, Austria) using a 2° cone and plate geometry. To facilitate cell attachment, the gels were functionalized with rat tail Collagen type I (0.2 mg/mL Corning, New York, New York, #354236) overnight at 4°C using the heterobifunctional crosslinker Sulfo-SANPAH (Thermo Scientific, Waltham, Massachusetts, 22,589; 0.4 mM in 50 mM HEPES buffer at pH 8) active with UV light (λ = 365 nm) for 10 min.

### Electron Microscopy

Cells attached to Aclar membrane (Agar Scientific) were fixed for 2 h in 2.5% glutaraldehyde (w/v) in 100 mM sodium cacodylate (CAB) buffer pH 7.2. Samples were washed twice for 10 min in 100 mM CAB and then post-fixed in 1% osmium tetroxide (w/v) in 100 mM CAB for 1 h before being dehydrated using an ethanol series of 50%, 70%, 90% (v/v) and 3 times with 100% ethanol for 10 minutes per step. The samples were then placed into propylene oxide, for (2x)10 min, and following this into a 1:1 mixture of propylene oxide and Agar LV resin (Agar Scientific) for 30 min. Following this, samples were embedded in freshly prepared Agar LV resin twice for 2 h before being placed in shallow aluminium moulds with the cells facing up and were polymerized at 60°C for 24 hours before being examined with a dissecting microscope to identify areas confluent with cells. These areas were cut out with a jig saw and attached to polymerized resin blocks with superglue, and once attached, the Aclar membrane was peeled off, exposing a monolayer of cells in the block face. Sections of 70 nm were cut on a Leica EM UC7 ultramicrotome using a diamond knife (Diatome) and were collected on 400 mesh copper grids. Sections were counterstained in 4.5% uranyl acetate (w/v) in 1% acetic acid (v/v) for 45 min and in Reynolds’ lead citrate for 7 min. Samples were viewed in a Jeol 1230 transmission electron microscope and images were captured with a Gatan One View 16mp camera.

### Data Availability

The data supporting the findings of this study are available from the corresponding author on request.

