## Supplementary Figures for "DNA damage alters nuclear mechanics through chromatin reorganisation"

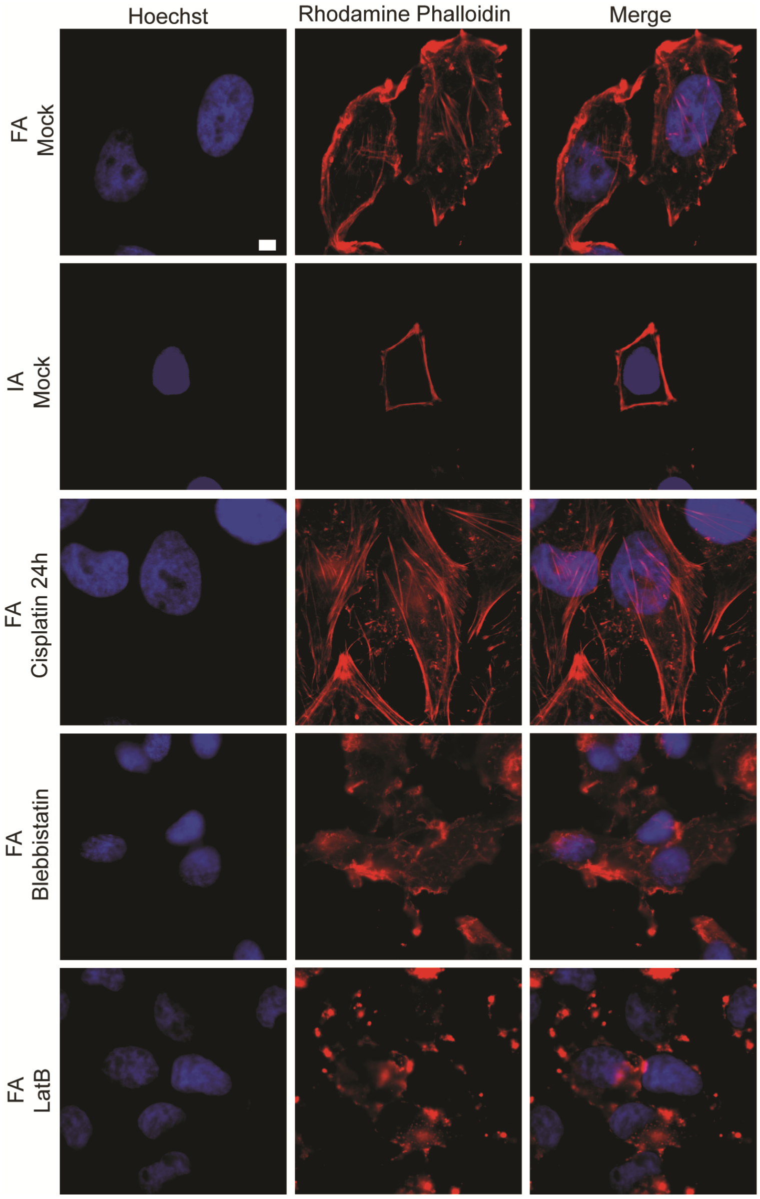


**Supplementary Figure 1 – Actin labelling of HeLa cells under different conditions.** Wide-field microscopy of actin labelled with Rhodamine Phalloidin, shown in red, and nuclear stain Hoechst, in blue, under stated conditions. (Scale bar = 5µm).

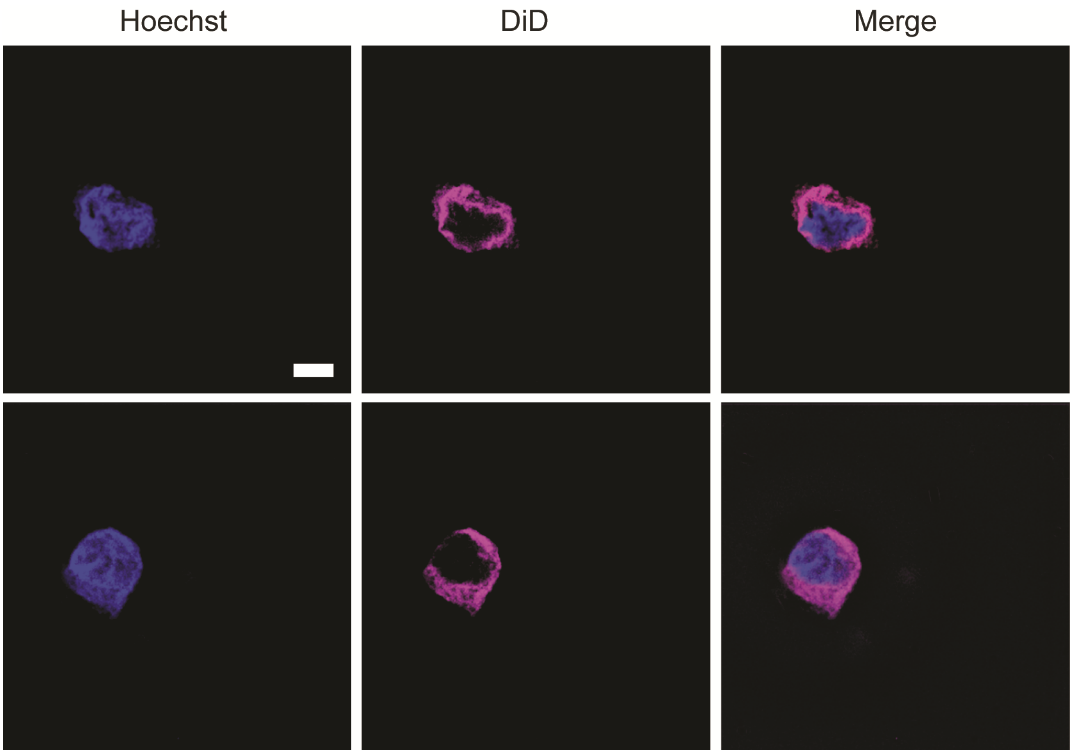


**Supplementary Figure 2 – Membrane integrity in isolated nuclei.** Wide-Field image of isolated nuclei with Hoechst DNA labelling and membrane staining using DiD. (Scale bar = 5μm).

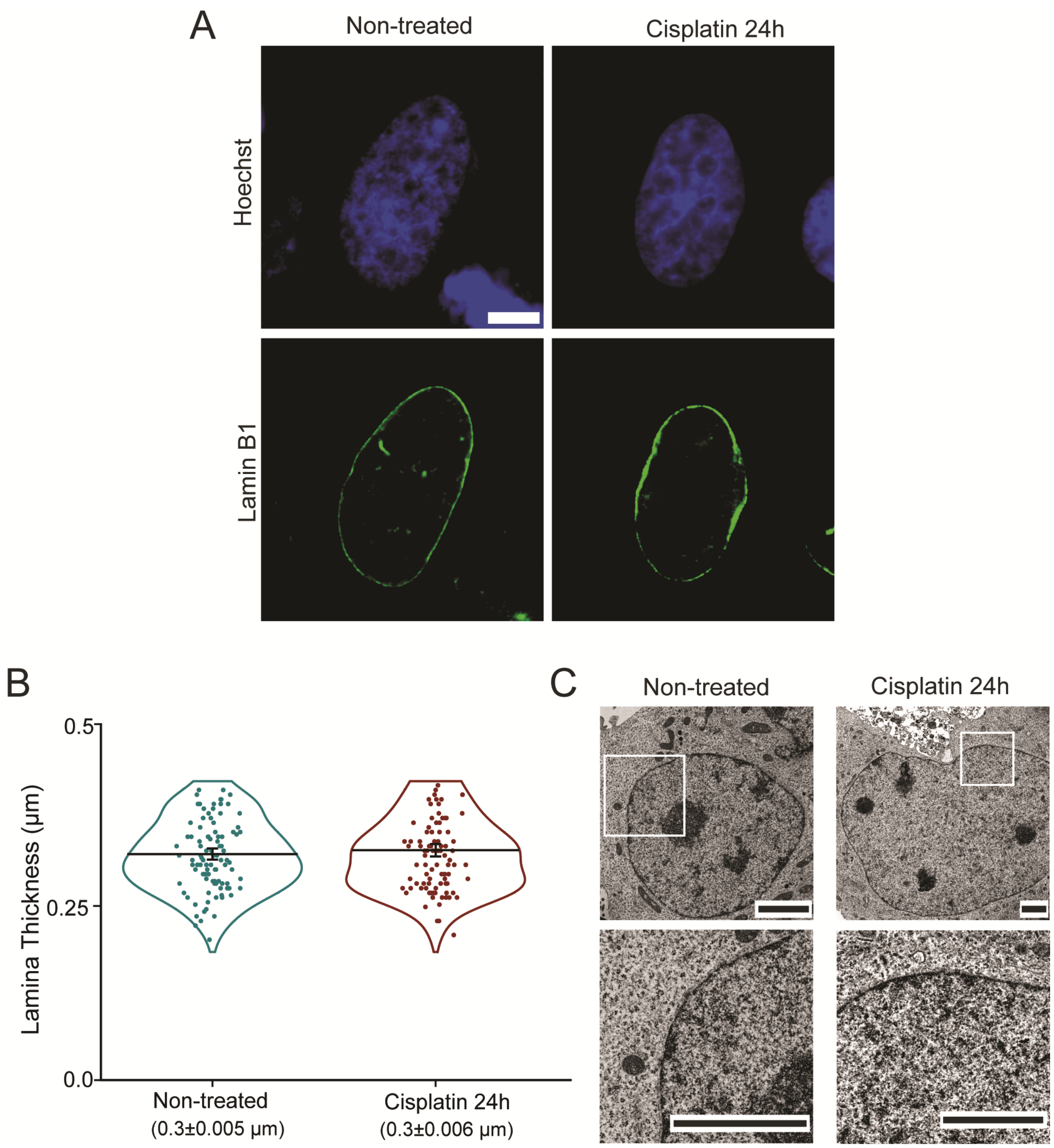


**Supplementary Figure 3 – Nuclear envelope integrity in HeLa cells after cisplatin treatment. (A)** Wide-field immunofluorescent images of Lamin B1 in non-treated and cisplatin-treated cells (green). (Scale bar = 5µm). **(B)** Measurements of nuclear lamina thickness, using Lamin B1 as a marker. Each point represents the average of 5 measurements at different regions in the nucleus of non-treated ($n$ = *101*) and long cisplatin treatment ($n$ = *100*) $p > 0.05$, calculated using a two-tailed t-test and assuming equal variance. Mean ± SE values are plotted. **(C)** Electron microscopy images showing membrane integrity in nuclei of both conditions. Lower panel shows zoomed in region in white square. (Scale bar = 2µm).
